## Supplementary figures and tables for "Aryl hydrocarbon receptor-dependent and -independent pathways mediate curcumin anti-aging effects"

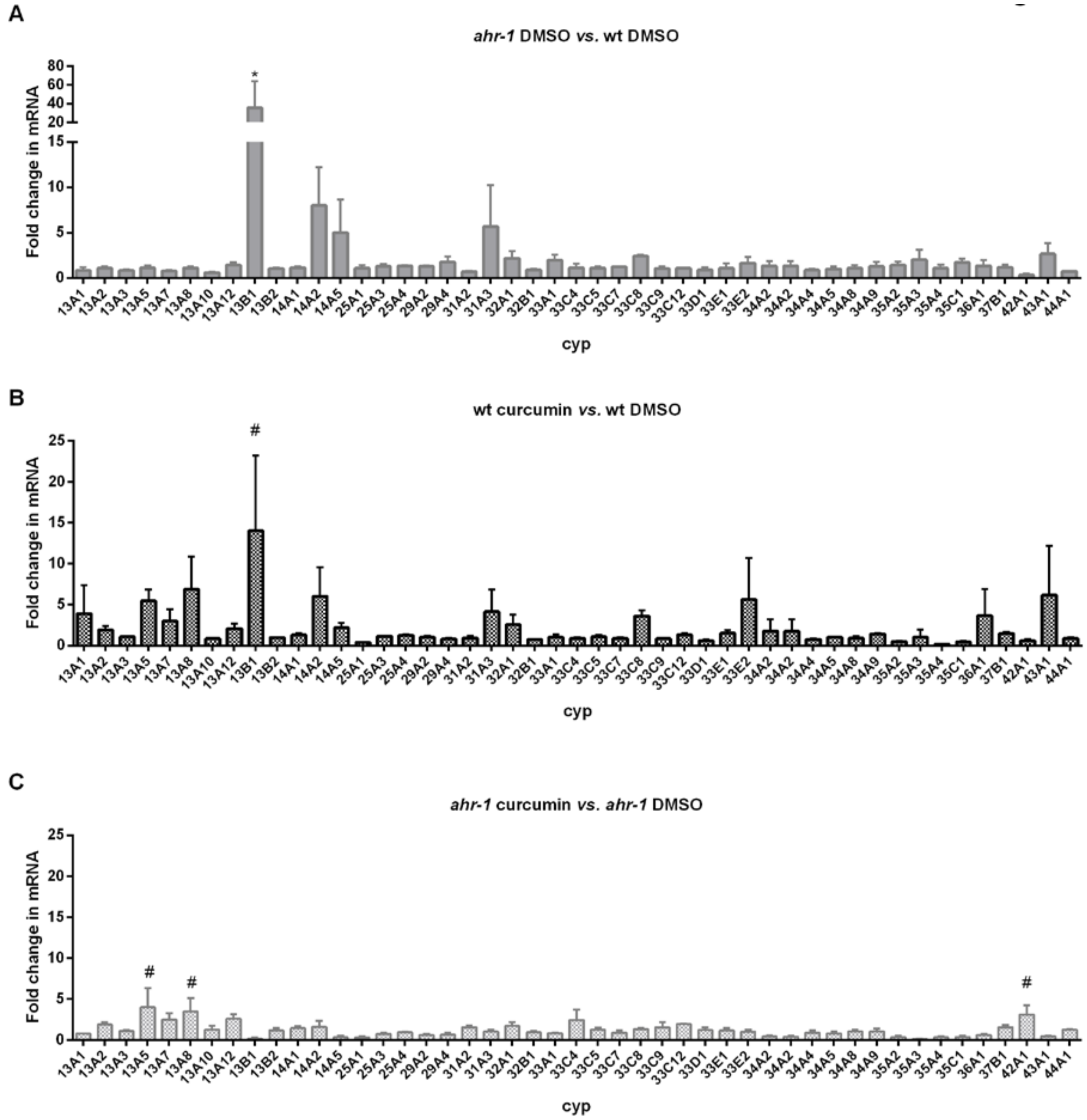


**Figure S1.** ***Cyp-13B1*** **expression is increased by curcumin in an** ***ahr-1*****-dependent manner.**

**A)** *cyp* gene expression in DMSO-treated *ahr-1(ju145)* relative to DMSO-treated wild-type. **B)** *cyp* gene expression in curcumin-treated wild-type relative to DMSO-treated wild-type. **C)** *cyp* gene expression in curcumin-treated *ahr-1* mutants relative to DMSO-treated *ahr-1* mutants. Mean + SEM of pooled data from 3 replicates is shown in all panels. * p-value < 0.05 *vs.* wt, # p-value < 0.05 *vs.* DMSO, statistical test: 2-way ANOVA with Tukey’s multiple comparisons test.

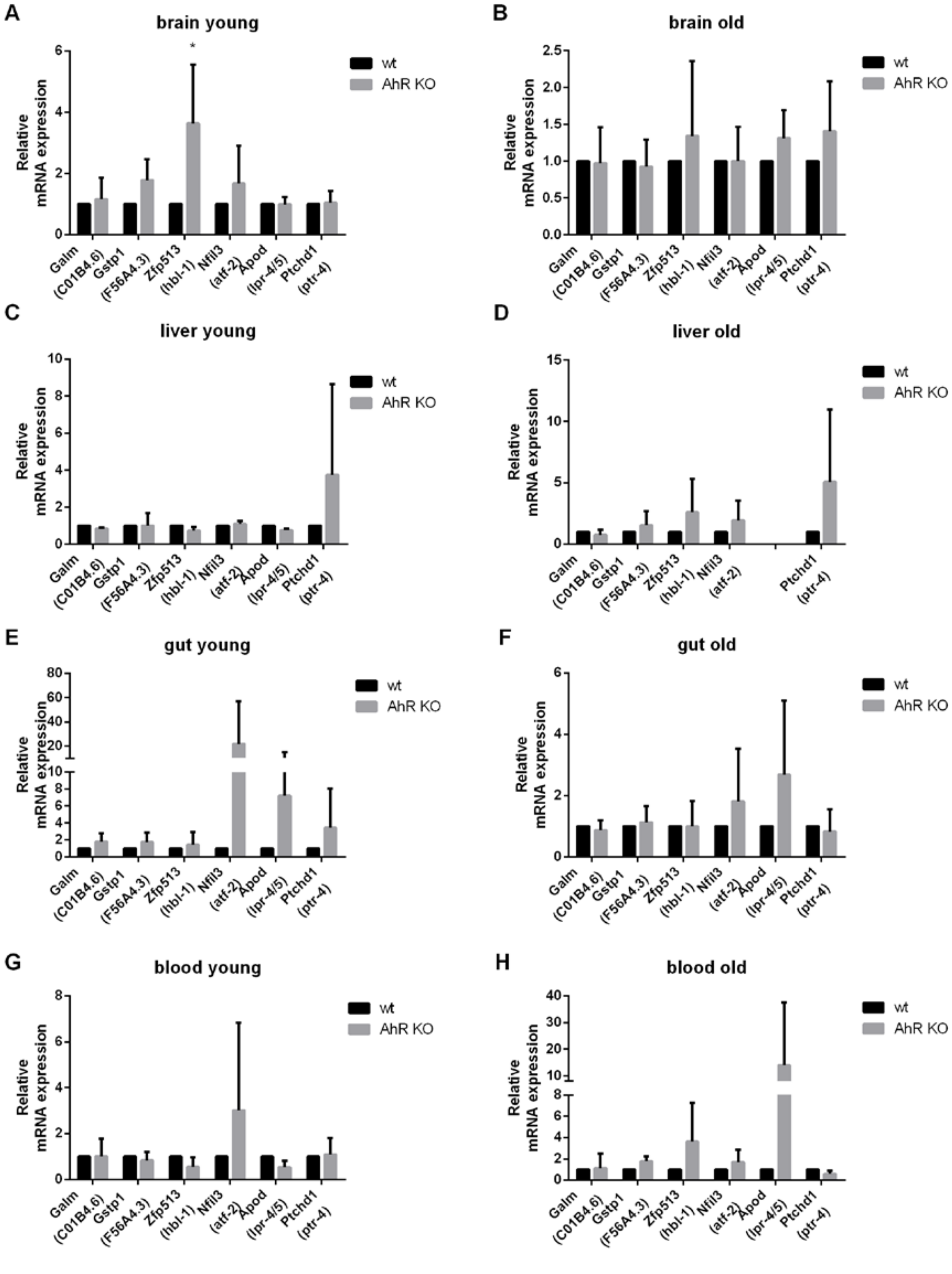


**Figure S2. Tissue-specific gene expression changes in mice.**

Expression levels of homologs of some of the strongest differentially expressed genes between wt and *ahr-1 C. elegans* in the brain **(A-B)**, liver **(C-D)**, gut **(E-F)**, and blood **(G-H)** of young (8-12 weeks old) or old (18 months old) mice was assessed by qPCR. The *C. elegans* homolog names are indicated in brackets. Mean + SD of 3 mice is shown. Statistical test: 2-way ANOVA with Sidak's multiple comparisons test, * p-value < 0.05 vs. wt.

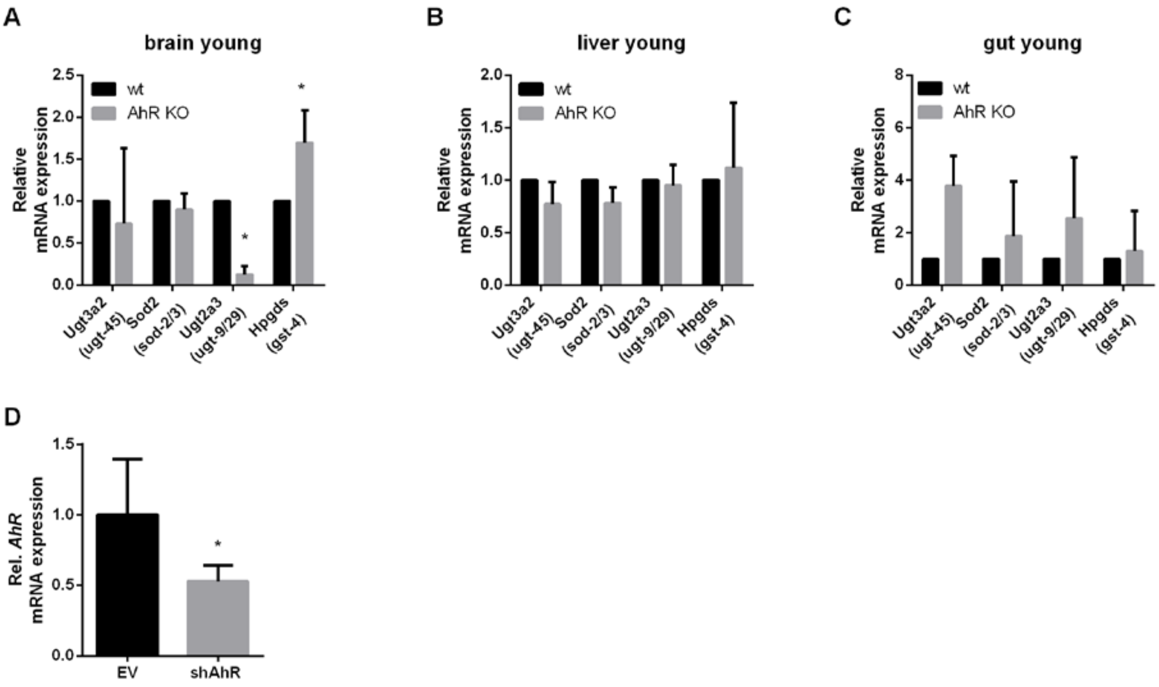


**Figure S3. Genes coding for detoxification enzymes are differentially expressed in a tissue-specific manner in mice.**

qPCR analysis of Ugt3A2, Sod2, Ugt2A3, and Hpdgs expression in the brain **(A)**, liver **(B)**, and gut **(C)** young (8-12 weeks old) or old (18 months old) mice. Mean + SD of pooled data from 3 mice are shown. Statistical test: 2-way ANOVA with Sidak's multiple comparisons test, * p-value < 0.05 vs. wt. **D)** Human primary EC were transfected with an empty vector (EV) or an expression vector for an shRNA targeting the human AhR transcript (shAhR). Relative *AhR* expression was assessed by qPCR, mean expression in the EV transfected cells was set to 1. Data are shown as mean + SD from 7 experiments. $^{\#}p<0.05$ vs. EV.

**Table S1. Primer pairs used for qPCR with *C. elegans* samples.**

| **Gene name** | **Forward Primer sequence (5' – 3')** | **Reverse Primer sequence (5' – 3')** | **Efficiency (%)[a]** |
|---|---|---|---|
| ***clec-209*** | TGCTCGGGGAACAACCAAAA | TTGGCTACGAACGATTGATGC | 88.4 |
| ***C01B4.6*** | TGGCGATGCGAAAATTGATGTAA | ATCTCCAGAAAGTGCTCGGC | 83.9 |
| ***F56A4.3*** | ACGAGGGGAATGAATGGCAA | CCATAGGGACCAATATCCATGAACT | 101.4 |
| ***C01B4.7*** | GTTTTGGAATCAGACGCGGG | CAGTGGGGTTCCGTCAAGTT | 106.6 |
| ***atf-2*** | CGAAGGAACAATGAAGCCGC | CCAAGAGCTGAACTCGTCGT | 98.1 |
| ***K04H4.2*** | ACGCCGGAATCTGTTGTTCT | CGTTCATTTGGAAAGGAGGCAT | 95.3 |
| ***egl-46*** | CACCTCAACCGCTTTTCCAAG | ATTTCACATCCGCCTCCTCC | 85.1 |
| ***T20F5.4*** | TCATCTACCGAGCAGCCAAC | GAGATGCTCGGTCTCACTGC | 94.5 |
| ***ptr-4*** | TCCTACCAGACGCGCAATC | GCAACCCATACTGACGGAGT | 79.5 |
| ***dyf-7*** | GTCTGCGTTTCCGTCACAAG | CGGGGAAGCAACAAGTTCTG | 124.8 |
| ***ugt-9*** | ATGTTGCCCAATAATTTGCT | TGGTTGGAAGACTGTAACAT | 100.2 |

| | | | |
|---|---|---|---|
| ***ugt-29*** | GATGGTGACTAAGGAAATCAAC | CCGATTTTACCAGTGATCCA | 102.8 |
| ***ugt-45*** | GAAATTCGGGTTCAGTCACA | TTAATAGCGTCGGTAGTCGG | 88.0 |
| ***ugt-57*** | CCTTGTCCGTGAGAAAGTTA | TCCGTCAATCCTTCCGTAT | 101.1 |
| ***cyp-13A1*** | GTACAACCTACACAAATCGC | TGAATCTCTGGATGAGTTGC | 91.2 |
| ***cyp-13A2*** | GTGGACAAAGAAATATGGCAA | ACAAGTGAACTCGTTGTTCT | 97.6 |
| ***cyp-13A3*** | GCCTGAAAGATGGGAATCTG | TGCCAGAAGCATCTTTTCTT | 100.5 |
| ***cyp-13A5*** | TGGACAAAAGAATACGGACC | CGTAGGTGATGACAGAGTTC | 86.7 |
| ***cyp-13A7*** | GACAAAGAAATACGGACCTG | TGCAGTTAATTTTCTCCCGT | 93.2 |
| ***cyp-13A8*** | AGCTCAAGGTTTTAGATGGA | CATAAGCTCCTGTGTGCTAT | 94.0 |
| ***cyp-13A10*** | AAGACAGTGCGATGGAATTA | TCTCCGAATATCGCTCTGA | 103.7 |
| ***cyp-13A12*** | CTGGGAGCTTTAGCAAATTC | TCTCCTGATTCCCATCTTTC | 105.8 |
| ***cyp-13B1*** | GGGCTGTTCCAGTGTTAGTC | AACATCGTGAACCTGCTTTA | 96.2 |
| ***cyp-13B2*** | TGAGAATGTATCCAGTTGCC | GATTCGAGCCATCTTTCTGG | 98.7 |
| ***cyp-14A1*** | TTCACCAGTTCCCTCCAGAC | AGAACGGTGAGCTCGGTAAG | 94.4 |
| ***cyp-14A2*** | AGTCCGGAAGGAAATACTTG | CATCTCGGTTTGTATGAGTC | 102.3 |
| ***cyp-14A5*** | ATAGGCACGGCGAGACTACC | CAAGTACAGTCTTGTTCAAC | 100.1 |
| ***cyp-25A1*** | TGGCTCTCCTGATTTTAAGC | AAGTTTTTCCTTGCGTTCTG | 84.7 |
| ***cyp-25A2*** | CTCTTCTGATTCTCACCTCAA | TTTTTCTTTCCGGTCAACGA | 93.1 |
| ***cyp-25A3*** | AGCACTTCTTTCCTTTTCCC | TAAACATCTGCAAGTCCCTC | 109.2 |
| ***cyp-25A4*** | CCAGTGTCTTTGATACTTCC | ACCCTCTCCAGCACGTCTTC | 106.4 |
| ***cyp-29A2*** | TGCCATAATTCTGGCATATC | AGGACCAGGTAGTTTACTTC | 90.3 |
| ***cyp-29A4*** | TTCCAGTGGCTCTTGCATTG | CATCTTTCCTCCATAAATCCAG | 107.4 |
| ***cyp-31A2*** | CCGCTGTACTTCTCGCTATG | TTCTTCATCTGCTCCGAGGC | 84.3 |
| ***cyp-31A3*** | TACTCCGCCGATCTCGTG | TTCTTCCTCTGCTCCGAGGG | 81.8 |
| ***cyp-32A1*** | TCCTAGCTGATGCAGTCGAG | CCGTTCTTGGATGACCTTTC | 94.6 |
| ***cyp-32B1*** | AGTTTTTGCATGTTCGGAAG | ACCATAACTTCAACACACCA | 97.8 |
| ***cyp-33A1*** | GAAGTTAATATTCAGGAACAT | TACGCGAAGTAGACACGAAG | 100.6 |
| ***cyp-33C4*** | AAAATCTTCCCAGACCCCT | GAAGACGGTGAGATCTTGTATCTA | 107.2 |
| ***cyp-33C5*** | GTTTGTTTGTTCCTGTTCCA | CATATGGAATGTTGCCCATC | 92.4 |
| ***cyp-33C7*** | GAAGATTTGATGAAAGACTGGA | CAGAGGTCCAAGCAAGTATT | 97.3 |
| ***cyp-33C8*** | GATGATGTGCTCAACTACTG | CTTGAGCCTTTTGCTTCTTC | 111.6 |
| ***cyp-33C9*** | ATTTTCAAGCCGGAGAGATT | CGGAACAACCCTGTATCTAT | 111.0 |
| ***cyp-33C12*** | ATATTGTCCCGATCAACCAG | TGGGTCTGGGAAAACCTTAT | 102.1 |
| ***cyp-33D1*** | CGTTTTGCAACAAACAACTC | CCAATGACTCTGTCTAGCTC | 106.1 |
| ***cyp-33E1*** | CAATGCAACTGTTAATGAATCT | TCAGGGAATATCTCTGGATC | 103.6 |

| | | | |
|---|---|---|---|
| ***cyp-33E2*** | ATGAATCACAACGTCTTGCC | TCAGGGAAGATTTCTGGATT | 100.2 |
| ***cyp-34A2*** | TGTGCGTGTCTTCTCAATAA | TTGGACTTCACTAATCACGG | 86.4 |
| ***cyp-34A4*** | TGAACGATTTGAACAGGGTG | TTCTCTGCACATTTCTTTGC | 81.4 |
| ***cyp-34A5*** | AGTTGTAGAGAAGCTGAGGA | GAACCTCATTAACAACCGCA | 98.9 |
| ***cyp-34A8*** | AGCTGTTTTTGATAACCGGAA | GGAGCAAATGGAGTAGTTGT | 109.3 |
| ***cyp-34A9*** | GGCGCAACTATCAATGAAATCC | TCAGCTCTAGCCAGTGATTC | 87.6 |
| ***cyp-35A2*** | TTCTGTGCTTTTGGGATACC | TATTACCGTACCTCTTTCTA | 79.6 |
| ***cyp-35A3*** | CGCTGCGTGTTTAAGTTGGC | TGTTGCCGTATTTCTTTCTG | 85.6 |
| ***cyp-35A4*** | TCGGCAATTTTGAGTTGGTT | TGTTGCCATATCTCTTTCTA | 107.7 |
| ***cyp-35C1*** | GAGCCGAGCTGTATTTAATC | ATGTCGAAGGCTTCGCATCT | 97.4 |
| ***cyp-36A1*** | GATCGATTCTTGAATAGTCGTG | TCGTGATGCTCGGACTGTAA | 79.8 |
| ***cyp-37B1*** | ATGTTTGAAGGCCACGACAC | TCCGGGTTGTACTGATTTGG | 89.8 |
| ***cyp-42A1*** | TCTACAACAGGCACGGAAGG | TGTGTCGTGGCCTTCAAATG | 122.2 |
| ***cyp-43A1*** | AGTCTGCTCGGATTCTTTTT | GTGCCGTAATTTCCTCATAG | 82.5 |
| ***cyp-44A1*** | ATCGGGAACATTGGGTATTT | CAGTTTGAACATCAGCAGGA | 81.5 |
| ***cdc-42*** | ATTACGCCGTCACAGTAATG | ATCCCTGAGATCGACTTGAG | 80.8-107.2 |
| ***act-1*** | GCTCTTGCCCCATCAACCAT | CACTTGCGGTGAACGATGGA | 94.8-100.4 |

[a] Primer efficiency is defined as how many copies of PCR product are produced per cycle. In ideal conditions, the primer efficiency is 100%, meaning that the amount of PCR product is doubled with each cycle. The primer efficiency was used for the calculation of the expression levels.

**Table S2. Primer pairs used for qPCR with mouse samples**.

| **Gene name** | **Forward Primer sequence (5' – 3')** | **Reverse Primer sequence (5' – 3')** | **Primer efficiency (%)[a]** |
|---|---|---|---|
| ***Ugt3a2*** | GTGTGTCGCAAGTTCTTCAT | TGATGTTTGGACCTGCCATA | 94.7 |
| ***Hpgds*** | CAGCGTTGGAGCAATGTCAA | CTGCCCAGGTTACATAATTGC | 109.9 |
| ***Ugt2a3*** | TATAGACCCTGTCGTTCCCTGT | TTGTGGGCCTTCCTAGGGTT | 92.6 |
| ***Sod2*** | AACGCCACCGAGGAGAAGTA | TCCAGCAACTCTCCTTTGGGT | 102.2 |
| ***Galm*** | AAGGAACTGCCTTCGACCTG | GACGCACCCTTGCACAAAAC | 111.0 |
| ***Gstp1*** | TGAGTACCCCTCTGTCTACGC | AGCTGCCCATACAGACAAGTG | 96.4 |
| ***Zfp513*** | GGGAGGGAGATTCCACAAGC | CCCCTGAGAGTCTCCTTCAGA | 146.9 |
| ***Nfil3*** | CAGGGAGCAGAACCACGATAAC | CCTCGTCCTACAGACCGGAT | 89.9 |
| ***Apod*** | ATCTGAGAGAAACTGACAGTGAGC | GAGACGGGCATTTCCCAAGA | 108.8 |
| ***Ptchd1*** | CCCTTTCGTCATGCTAGGTCAT | GCCGAAGGTGACCAAGTACA | 123.2 |

[a] Primer efficiency is defined as how many copies of PCR product are produced per cycle. In ideal conditions, the primer efficiency is 100%, meaning that the amount of PCR product is doubled with each cycle.
